# Genetic Divergence Analysis among Sesame (*Sesamum Indicum* L.) Accessions in Middle Awash, Ethiopia

**DOI:** 10.1101/2022.12.29.522196

**Authors:** Mesay Tadesse Gimbichu, Gudeta Nepir Gurmu, Negash Geleta Ayana

**Affiliations:** Ethiopian Institute of Agricultural Research, Werer Agricultural Research Center, P.O. Box: 2003, Addis Ababa, Ethiopia.; Ambo University College of Agriculture and Veterinary Science, Department of Plant Science, P.O.Box.19, Ambo, Ethiopia.; Ethiopian Institute of Agricultural Research, Kulumsa Agricultural Research Center, P.O. Box: 2003, Addis Ababa, Ethiopia.

**Keywords:** Accessions, Cluster analysis, genetic distance, genetic divergence, principal component, sesame

## Abstract

This research was conducted to assess genetic distance, extent, and pattern of diversity among sesame accessions. A total of 64 sesame Accessions were evaluated in an 8 x 8 lattice design with two replications in 2021 at Werer Agricultural Research Center. Analysis of variance revealed that there was a statistically significant difference among the accessions for all traits except for 50% days to emergence and the number of seeds per pod. Principal components analysis showed the first five PCAs viz. PC1 (21.9%), PC2 (11.00%), PC3 (15.6%), PC4 (18.3%), and PC5 (9.5) with a total contribution of 76.3% variation. The dendrogram was constructed using the Unweighted Pair-group Method with Arithmetic Means to separate Accessions into five distinct clusters. Sesame accessions with high seed yield and high mean values for other desirable traits were grouped into Cluster I Cluster V. Cluster IV and Cluster V had the highest inter-cluster distance. Accession in Cluster V (Acc.241297) could be crossed with other clusters to come up with promising segregation for further improvement programs.

## INTRODUCTION

Sesame (*Sesamum indicum* L.) is the most important oil crop grown well in tropical and sub-tropical climates (Daniel and Parzies, 2011). Cultivated sesame belongs to the order Tubiflorae and the family Pedaliaceae, with 37 species described under the genus Sesamum, of which only (*Sesamum indicum* L) has been recognized as a cultivated species (Getinet *et al*., 1997). The cultivated species sesame (*Sesamum indicum* L.) is a diploid species with a chromosome number of 2n = 2x = 26 (Morinaga *et al*., 1929).

According to Getinet *et al*. (1997), Ethiopia is considered to be the center or origin of sesame and it seems that the crop was taken into cultivation in other African countries and then taken to India. The presence of weedy or wild forms of sesame (S. alatum; 2n = 26 and S. latifolium, 2n = 32) in Ethiopia shows that it is indigenous and considered the center of origin for sesame. Also, the presence of high genetic diversity is serving as a resource for further improvement of the crop (Daniel and Parzies, 2011).

Sesame, locally called *‘Selit’*, is a valuable export crop in Ethiopia. Ethiopia is the world’s fourth-largest exporter of sesame seeds, after Sudan, India, and Nigeria, and Africa’s third-largest exporter, after Sudan and Nigeria (FAOSTAT, 2020). Ethiopian Central Statistics Agency reported the area under production of sesame is estimated to be 294,819.49, 375,119.95, and 369,897.32 ha in the 2018/2019, 2019/2020, and 2020/2021 cropping seasons, respectively (CSA, 2019; CSA, 2020; CSA, 2021). According to CSA, the total production of sesame was 262, 665.4 and 260, 257.6tonha-1, in 2019/2020, and 2020/2021 cropping seasons, respectively (CSA, 2020; CSA, 2021). Total grain production and productivity are 260, 257.6 ton ha-1, and 700 kg ha-1, respectively during 2019/2020 and 2020/2021 cropping seasons (CSA, 2021).

The knowledge of diversity and genetic distances of genotypes helps to identify parental lines for hybridization programs, which can be used to select appropriate parental genotypes for hybridization to develop a high-yielding potential variety (Bhatt, 1970). Principal component analysis (PCA) is the most essential tool in diversity studies. This technique is highly effective and useful for identifying plant features that classify the distinctiveness of promising genotypes (Ashim *et al*., 2013). Shammoro *et al*. (2020) also reported parents chosen from divergent clusters are expected to have the most genetic recombination. To select the desired parent for crossing program and to develop new crop varieties; understanding and knowing genetic diversity is the most crucial in plant breeding. For that reason, the present research was undertaken with the objective to assess genetic distance and clustering among sesame accessions.

## MATERIALS AND METHODS

### Description of Experimental Site

The experiment was conducted during 2021 in the summer season at Werer Agricultural Research Center (WARC) in the low land of Ethiopia (middle awash) of Afar Regional State under irrigation condition. Werer is located at a latitude of 9° 16’ N, the longitude of 40° 9’ E, and an altitude of 740 m a.s.l. The mean annual rainfall of the area is 590 mm and the mean annual minimum and maximum temperature ranges between 26.7 and 40.8°C, respectively (www.eiar.gov.et).

### Experimental Materials

The experimental materials consisted of 64 sesame accessions obtained from Ethiopian Biodiversity Institute (EBI) were used in this study (Table 1). The origin of those materials was Ethiopian country and which was collected from four regions (Benshungul Gumuz, Tigray, Oromia, and Amhara).

**Table 1.**
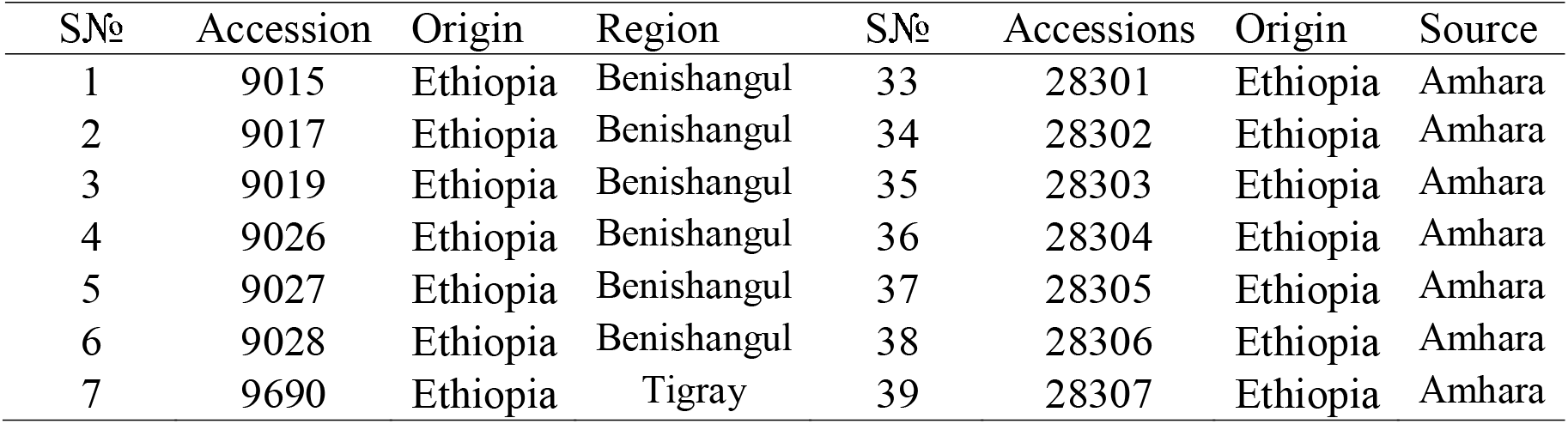
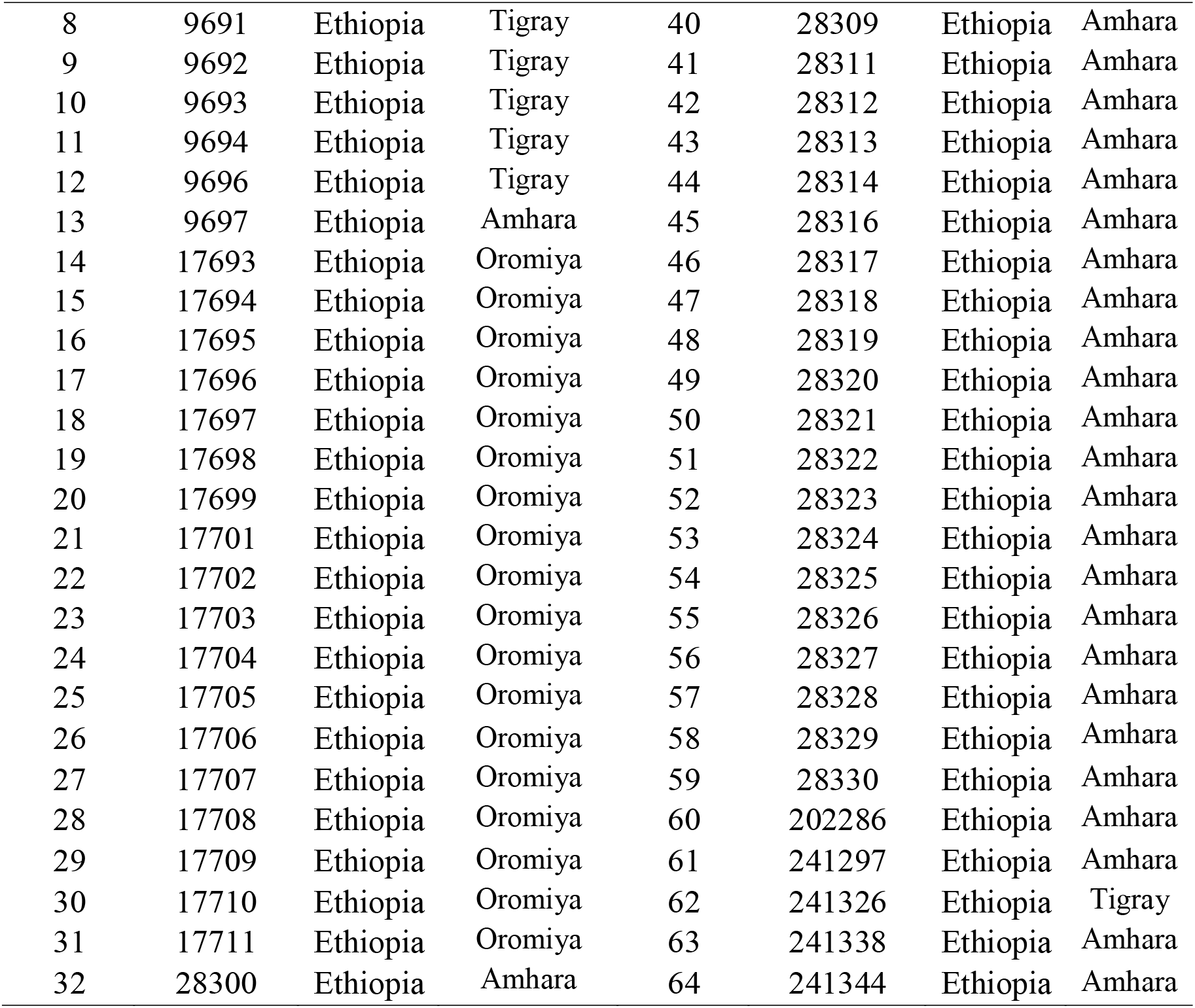
Experimental materials used for the study

| SN<sub>o</sub> | Accession | Origin | Region | SN<sub>o</sub> | Accessions | Origin | Source |
| --- | --- | --- | --- | --- | --- | --- | --- |
| 1 | 9015 | Ethiopia | Benishangul | 33 | 28301 | Ethiopia | Amhara |
| 2 | 9017 | Ethiopia | Benishangul | 34 | 28302 | Ethiopia | Amhara |
| 3 | 9019 | Ethiopia | Benishangul | 35 | 28303 | Ethiopia | Amhara |
| 4 | 9026 | Ethiopia | Benishangul | 36 | 28304 | Ethiopia | Amhara |
| 5 | 9027 | Ethiopia | Benishangul | 37 | 28305 | Ethiopia | Amhara |
| 6 | 9028 | Ethiopia | Benishangul | 38 | 28306 | Ethiopia | Amhara |
| 7 | 9690 | Ethiopia | Tigray | 39 | 28307 | Ethiopia | Amhara |
| 8 | 9691 | Ethiopia | Tigray | 40 | 28309 | Ethiopia | Amhara |
| 9 | 9692 | Ethiopia | Tigray | 41 | 28311 | Ethiopia | Amhara |
| 10 | 9693 | Ethiopia | Tigray | 42 | 28312 | Ethiopia | Amhara |
| 11 | 9694 | Ethiopia | Tigray | 43 | 28313 | Ethiopia | Amhara |
| 12 | 9696 | Ethiopia | Tigray | 44 | 28314 | Ethiopia | Amhara |
| 13 | 9697 | Ethiopia | Amhara | 45 | 28316 | Ethiopia | Amhara |
| 14 | 17693 | Ethiopia | Oromiya | 46 | 28317 | Ethiopia | Amhara |
| 15 | 17694 | Ethiopia | Oromiya | 47 | 28318 | Ethiopia | Amhara |
| 16 | 17695 | Ethiopia | Oromiya | 48 | 28319 | Ethiopia | Amhara |
| 17 | 17696 | Ethiopia | Oromiya | 49 | 28320 | Ethiopia | Amhara |
| 18 | 17697 | Ethiopia | Oromiya | 50 | 28321 | Ethiopia | Amhara |
| 19 | 17698 | Ethiopia | Oromiya | 51 | 28322 | Ethiopia | Amhara |
| 20 | 17699 | Ethiopia | Oromiya | 52 | 28323 | Ethiopia | Amhara |
| 21 | 17701 | Ethiopia | Oromiya | 53 | 28324 | Ethiopia | Amhara |
| 22 | 17702 | Ethiopia | Oromiya | 54 | 28325 | Ethiopia | Amhara |
| 23 | 17703 | Ethiopia | Oromiya | 55 | 28326 | Ethiopia | Amhara |
| 24 | 17704 | Ethiopia | Oromiya | 56 | 28327 | Ethiopia | Amhara |
| 25 | 17705 | Ethiopia | Oromiya | 57 | 28328 | Ethiopia | Amhara |
| 26 | 17706 | Ethiopia | Oromiya | 58 | 28329 | Ethiopia | Amhara |
| 27 | 17707 | Ethiopia | Oromiya | 59 | 28330 | Ethiopia | Amhara |
| 28 | 17708 | Ethiopia | Oromiya | 60 | 202286 | Ethiopia | Amhara |
| 29 | 17709 | Ethiopia | Oromiya | 61 | 241297 | Ethiopia | Amhara |
| 30 | 17710 | Ethiopia | Oromiya | 62 | 241326 | Ethiopia | Tigray |
| 31 | 17711 | Ethiopia | Oromiya | 63 | 241338 | Ethiopia | Amhara |
| 32 | 28300 | Ethiopia | Amhara | 64 | 241344 | Ethiopia | Amhara |

### Experimental Design and Field Management

The experiment was planted in an 8 × 8 simple lattice design. The spacing between plants and rows was 10 cm and 40 cm respectively. Then the gross plot size was 2.4m * 3m=7.2m2 and net plot size was 4.8m2 (four rows). According to recommended rate; the seed rate used for planting was 5-7kg/ha which means on average 4 g was required per plot. The plant population was adjusted by thinning after germination according to recommended pant per hectare (25, 0000/ha) and 180 plants per plot. After seedlings reached 15-20cm, thinning was made to adjust the distances of 10cm between plants. Crop management practices such as weeding, thinning, crop protection, and securing it from higher animal and related management which used for a crop was done regularly.

### Data collection

To avoid border effects, the data for all of the parameters considered were taken from the net plot area of 1.6 m x 3 m (4.8 m^2^). Data were collected for the following parameters using IPGRI sesame descriptor (IPGRI, 2004) for Days to 50% emergency (DE), Days to 50% flowering (DF), Days to 75% maturity (DM), Plant height (PH) (cm), Length of pod bearing zone (LPBZ) (cm), Number of primary branches per plant (PBP), Secondary branches per plant (SBP), Number of pods per plant (NPP), Pod length (CL) (cm), Pod width (CW) (cm), Biomass yield per hectare (BM) (Kg/ha), Seed yield (SY) (kg/ha), Thousand seed weight (TSW) (g), Harvest Index (HI) (%) and Oil content (OC): oil content.

### Data Analysis

#### Analysis of variance

The data were analyzed following procedures appropriate to the simple lattice design used as described by Gomez and Gomez (1984). Means of significant treatment effects were separated using Duncan’s New Multiple Range Test (DNMRT) test at a 5% probability level. Data collected for each character was subjected to analysis of variance using SAS software version 9.3 (SAS institute, 2014).

#### Principal Component Analysis

Principal component analysis (PCA) was computed to identify the characters which accounted more for the total variation. The data was standardized to mean zero and variance of one before computing principal component analysis. The principal component based on the correlation matrix was calculated using R- software.

#### Genetic Distance and Clustering

The genetic distance of 64 sesame accessions was estimated using clustering of genotypes. Euclidean distance (ED) was calculated from quantitative traits after standardization (subtracting the mean value and dividing it by the standard deviation) as established by Sneath and Sokal (1973) using Minitab v-17 software as follows:

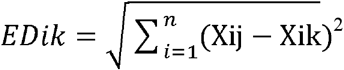

Where, EDjk = distance between genotypes j and k; xij and xik = phenotype traits values of the i^th^ character for genotypes j and k, respectively; and n = number of phenotype traits used to calculate the distance. The distance matrix from phenotype traits was used to construct a dendrogram based on the unweighted Pair-group Method with Arithmetic Means (UPGMA). The results of cluster analysis were presented in the form of a dendrogram.

## RESULTS AND DISCUSSION

### Analysis of variance

Analysis of variance (ANOVA) for 16 traits revealed that majority of the traits, such as days to 50% flowering, number of primary branches per plant, number of secondary branches per plant, number of pods per plant, pod length, pod width, biomass yield, seed yield, harvest index, thousand seed weight, days to 75% maturity, and oil content showed highly significant (*P*<0.01) variation and length of pod bearing zone and plant height was also significant at (*P*<0.05) (Table 2). This indicates the presence of genotypic variation among the tested sesame accessions. Accessions had recorded a non-significant difference for days to 50% emergence and the number of seeds per pod.

**Table 2.** Mean squares from analysis of variance for 16 traits of 64 sesame accessions

| SN. | Trait | Rep (Df=1) | Block (Df=14) | Genotype (Df=63) | Error (Df=49) | CV(%) |
| --- | --- | --- | --- | --- | --- | --- |
| 1 | Days to 50% emergence | 0.28<sup>ns</sup> | 0.36<sup>*</sup> | 0.204<sup>ns</sup> | 0.1557 | 7.03 |
| 2 | Days to 50% flowering | 164.257<sup>**</sup> | 46.237<sup>**</sup> | 32.73<sup>**</sup> | 11.69 | 5.64 |
| 3 | Days to 75% maturity | 29.07<sup>ns</sup> | 17.45<sup>ns</sup> | 96.619<sup>**</sup> | 14.348 | 3.33 |
| 4 | Plant height (cm) | 165.24<sup>ns</sup> | 224.46<sup>ns</sup> | 351.96<sup>*</sup> | 145.027 | 10.9 |
| 5 | Length of pod bearing zone(cm) | 1316.49<sup>**</sup> | 52.81<sup>ns</sup> | 82.59<sup>*</sup> | 36.61 | 11.6 |
| 6 | Primary branch plant<sup>-1</sup> | 0.06<sup>ns</sup> | 0.02<sup>ns</sup> | 0.279<sup>**</sup> | 0.014 | 4.15 |
| 7 | Secondary branch plant<sup>-1</sup> | 0.00397<sup>ns</sup> | 0.000<sup>ns</sup> | 0.008<sup>**</sup> | 0.0027 | 18.8 |
| 8 | Number of pod plant<sup>-1</sup> | 1774.59<sup>**</sup> | 25.71<sup>ns</sup> | 358.06<sup>**</sup> | 41.36124 | 14.1 |
| 9 | Pod length (cm) | 0.0009<sup>ns</sup> | 0.0004<sup>ns</sup> | 0.098<sup>**</sup> | 0.0005 | 0.84 |
| 10 | Pod width (cm) | 0.000078<sup>ns</sup> | 0.0014<sup>ns</sup> | 0.0034<sup>**</sup> | 0.00097 | 4.3 |
| 11 | Number of seed pod<sup>-1</sup> | 111.079<sup>ns</sup> | 37.056<sup>ns</sup> | 33.44<sup>ns</sup> | 31.8565 | 8.99 |
| 12 | 1000 seed weight (g) | 0.1069<sup>*</sup> | 0.0056<sup>ns</sup> | 0.273<sup>**</sup> | 0.0073 | 2.93 |
| 13 | Biomass yield (kg/ha) | 213170.22<sup>ns</sup> | 144248.43<sup>ns</sup> | 1433363.6<sup>**</sup> | 155152.9 | 10.0 |
| 14 | Seed yield (kg/ha) | 17543.689<sup>ns</sup> | 9777.53<sup>ns</sup> | 87466.8<sup>**</sup> | 11750.05 | 13 |
| 15 | Harvest index (%) | 3.78<sup>ns</sup> | 3.95<sup>ns</sup> | 16.45<sup>**</sup> | 3.56 | 8.94 |
| 16 | Oil content (%) | 23.69<sup>**</sup> | 3.145<sup>*</sup> | 4.02<sup>**</sup> | 0.798 | 1.81 |

\*\* And \*indicates highly significant at (1%), significant at (5%) probability levels respectively, ns= non-significant, CV= coefficient of variations, Df= degrees of freedom.

In line with the present findings, different researchers reported the presence of significant variation among sesame genotypes. For instance; Shammoro *et al*. (2020) and Tesfaye *et al*. (2021) reported highly significant differences among genotypes for days to 50% flowering, number of primary branches per plant, number of pods per plant, seed yield, thousand seed weight, and days to 75% maturity among 100 genotypes and 300 genotypes, respectively.

#### Principal Component Analysis

Principal component analysis is the most essential tool in diversity studies. The principal component analysis (PCA) revealed total five principal components (PCs) with Eigen values greater than one (ranging from 1.2 to 4.5) for 14 quantitative traits among the 64 sesame accession (Table 3). The five principal components accounted for percentages of total variance that ranged from 9.5 to 21.9% and accounted for 76.3% of the total variation. According to Brejda *et al*. (2000), data were considered in each component with an Eigenvalue of >1, which determined at least 10% of the variation. The higher Eigenvalues were considered the best representatives of system attributes in the principal component.

**Table 3.**
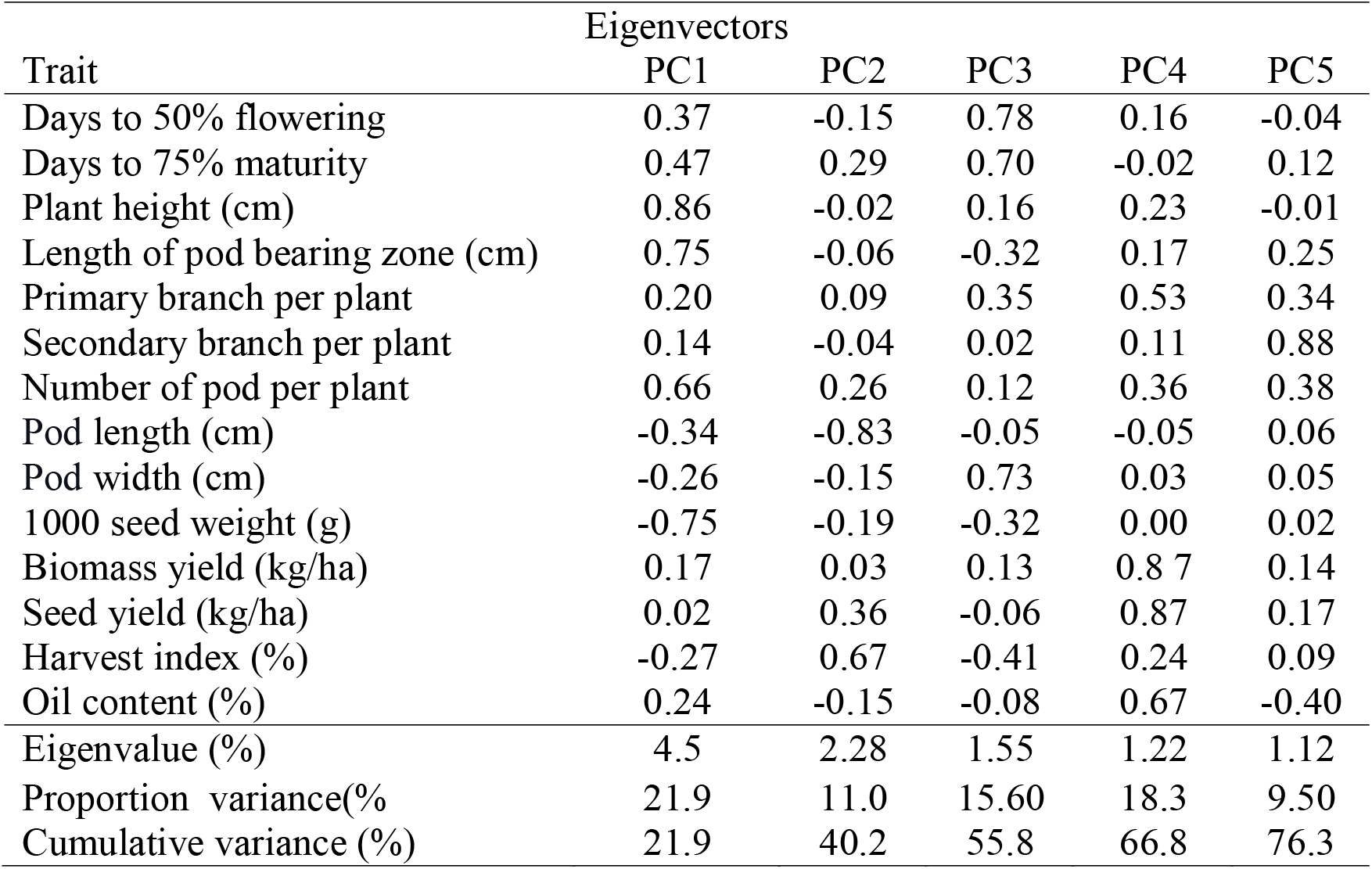
Principal components and Eigen values of the first five principal components for 14 quantitative traits of 64 sesame accessions.

| Trait | Eigenvectors |  |  |  |  |
| --- | --- | --- | --- | --- | --- |
|  | PC1 | PC2 | PC3 | PC4 | PC5 |
| Days to 50% flowering | 0.37 | -0.15 | 0.78 | 0.16 | -0.04 |
| Days to 75% maturity | 0.47 | 0.29 | 0.70 | -0.02 | 0.12 |
| Plant height (cm) | 0.86 | -0.02 | 0.16 | 0.23 | -0.01 |
| Length of pod bearing zone (cm) | 0.75 | -0.06 | -0.32 | 0.17 | 0.25 |
| Primary branch per plant | 0.20 | 0.09 | 0.35 | 0.53 | 0.34 |
| Secondary branch per plant | 0.14 | -0.04 | 0.02 | 0.11 | 0.88 |
| Number of pod per plant | 0.66 | 0.26 | 0.12 | 0.36 | 0.38 |
| Pod length (cm) | -0.34 | -0.83 | -0.05 | -0.05 | 0.06 |
| Pod width (cm) | -0.26 | -0.15 | 0.73 | 0.03 | 0.05 |
| 1000 seed weight (g) | -0.75 | -0.19 | -0.32 | 0.00 | 0.02 |
| Biomass yield (kg/ha) | 0.17 | 0.03 | 0.13 | 0.87 | 0.14 |
| Seed yield (kg/ha) | 0.02 | 0.36 | -0.06 | 0.87 | 0.17 |
| Harvest index (%) | -0.27 | 0.67 | -0.41 | 0.24 | 0.09 |
| Oil content (%) | 0.24 | -0.15 | -0.08 | 0.67 | -0.40 |
| Eigenvalue (%) | 4.5 | 2.28 | 1.55 | 1.22 | 1.12 |
| Proportion variance(%) | 21.9 | 11.0 | 15.60 | 18.3 | 9.50 |
| Cumulative variance (%) | 21.9 | 40.2 | 55.8 | 66.8 | 76.3 |

For the present results principal component one (PC1) had a relatively higher value and included the traits namely days to 50% flowering, plant height, number of pods per plant, length of pod bearing zone, and days to 75% maturity. These parameters made a greater impact on total diversity and were responsible for cluster differentiation. Traits like harvest index and seed yield contributed more to total genetic diversity in the principal component (PC2). In principal component three (PC3), the number of primary branches per plant, days to 50% flowering, days to 75% maturity and pod width had relatively greater contributions to the total genetic diversity. The number of primary branches per plant, number of pods per plant, biomass yield, percent oil content, and seed yield all contributed significantly to the total variance in principal component four (PC4). In the principal component five (PC5), characters such as the number of primary branches per plant, number of pods per plant, and the number of secondary branches per plant had relatively high contributions to the total variance.

### Genetic Divergence Analysis

#### Clustering of accession

Based on the accession mean values for 14 quantitative traits, the 64 sesame accessions were divided into five distinct clusters (Figure 1). Cluster II consisted of 25 accessions followed by Cluster I which consisted of 15 accessions, Cluster III had 13 genotypes, Cluster IV had ten accessions and Cluster V had one genotype. This finding indicated that accession belonging to the same cluster shared many traits which resemble each other but vary from accession belonging to other clusters in one or more traits. Similarly, Shammoro *et al*. (2020); Tesfaye *et al*. (2021); and Sultana *et al*. (2019) reported, seven, six, and five clusters for one hundred, three hundred, two hundred eighty-four, fifty genotypes respectively.

**Figure 1.**
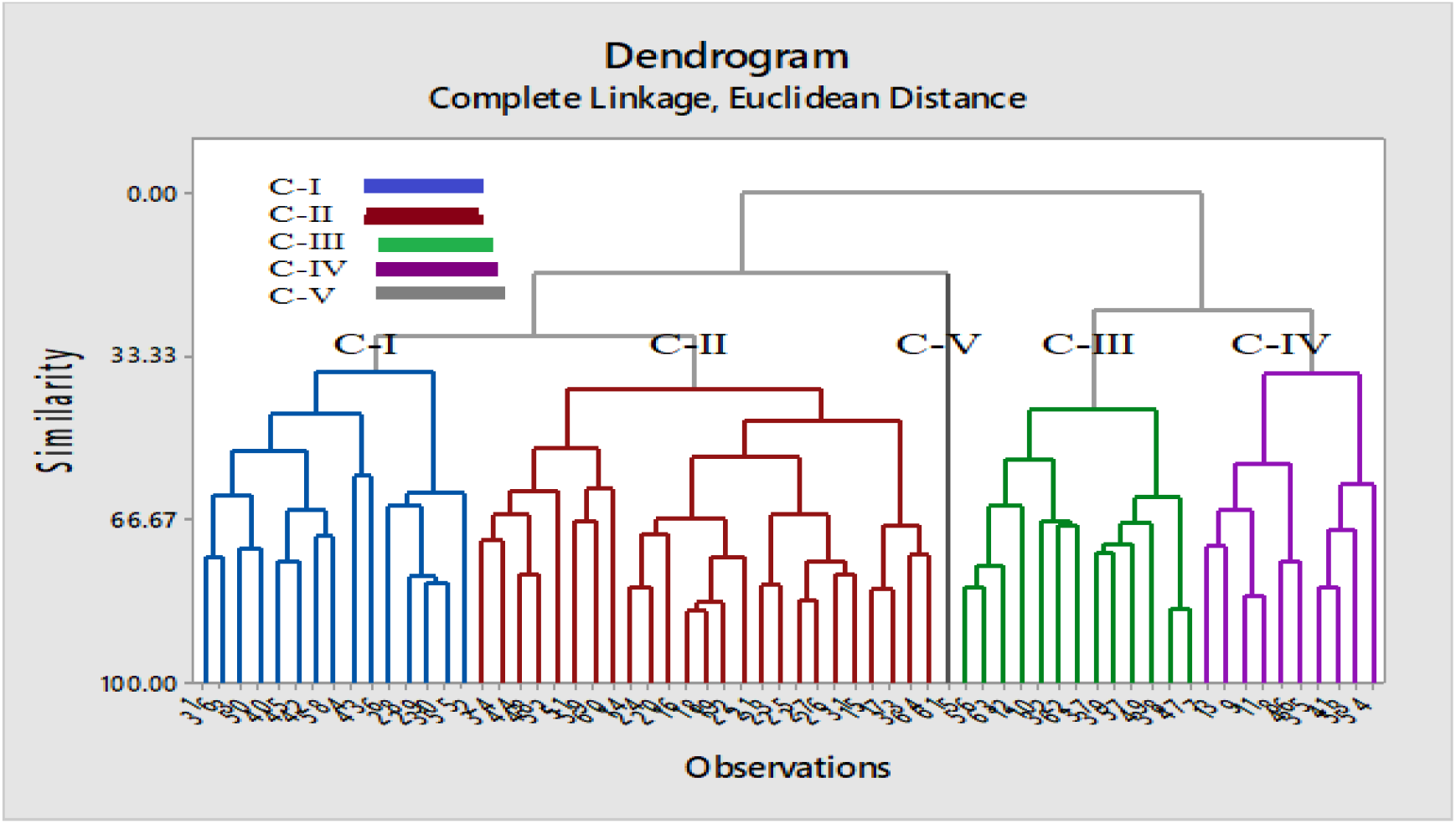
Dendrogram generated based on UPGMA clustering method depicting genetic relationships among 64 sesame accessions based on 14 quantitative traits.

#### Intra and Inter cluster distance of accessions

The intra-cluster and inter-cluster distances of 64 accession were listed in the Table 4, with Cluster IV and Cluster V having the highest inter-cluster distance, followed by Cluster III and Cluster V. Cluster V and the other clusters (clusters II, III, and IV) had a ranged from (6.45-8.88) an ED. Cluster V had a high dissimilarity level with all the other clusters. This means that the accession included in cluster V could be used as a parent during hybridization for the rest accession included in the other four clusters. The intercluster distance between Cluster I and Cluster II was the smallest of all (2.49). Because the accession in this group had a high similarity level or a low dissimilarity level, the effectiveness of hybridization and segregation will be low. Present result revealed that intra-cluster distances were lower than inter-cluster distances. This suggests that wider diversity among the accession of different groups and accession within the same cluster were closely related. Similar results were also reported by Sultana *et al*. (2019). The intra-cluster distances had a ranged from (0 to 2.95) an ED. High intra cluster distance was recorded for cluster IV followed by Cluster I, cluster II, cluster III and cluster V.

**Table 4.** Average intra (bold) and inter (off) diagonal cluster distance among 64 sesame accessions

| Cluster | Cluster-I | Cluster-II | Cluster-III | Cluster-IV | Cluster-V |
| --- | --- | --- | --- | --- | --- |
| Cluster-I | **2.92** | 2.49 | 4.51 | 4.31 | 6.45 |
| Cluster-II |  | **2.79** | 3.37 | 4.51 | 6.61 |
| Cluster-III |  |  | **2.75** | 3.4 | 8.75 |
| Cluster-IV |  |  |  | **2.95** | 8.88 |
| Cluster-V |  |  |  |  | **0** |

##### Cluster Mean Analysis

The mean values of the 5 clusters for 14 quantitative traits are presented in Table 5. The distinguishing features of Cluster I have higher mean values than overall mean values of accession for all traits except for 1000 seed weight and pod length. Also, Cluster V had higher mean values than the overall mean values of accession for all traits including for seed yield except for 1000 seed weight and harvest index. Cluster II had higher mean values for pod length, pod width, and 1000 seed weight. Cluster III had a high mean value for five traits, viz., and days to 50% flowering, plant height, primary branch per plant, oil content, and days to 75% maturity. Cluster IV had a high mean value for three traits namely plant height, length of pod bearing zone, and harvest index.

**Table 5.** Cluster mean values for 14 quantitative traits of 64 sesame accessions

| Trait | Cluster-I | Cluster-II | Cluster-III | Cluster-IV | Cluster-V | Grand mean |
| --- | --- | --- | --- | --- | --- | --- |
| DF | 61.73 | 58.85 | 61.63 | 59.69 | 61.23 | 60.62 |
| DM | 114.77 | 110.12 | 114.88 | 113.15 | 115.23 | 113.60 |
| PH | 117.57 | 98.06 | 112.38 | 113.47 | 114.03 | 110.87 |
| LPBZ | 54.94 | 49.37 | 50.25 | 52.43 | 55.83 | 52.28 |
| PBP | 3.05 | 2.53 | 2.92 | 2.61 | 2.91 | 2.80 |
| SBP | 0.30 | 0.27 | 0.26 | 0.28 | 0.29 | 0.28 |
| NPP | 55.13 | 35.70 | 43.40 | 41.85 | 55.27 | 45.58 |
| CL | 2.73 | 2.91 | 2.76 | 2.65 | 2.75 | 2.76 |
| CW | 0.74 | 0.73 | 0.71 | 0.72 | 0.72 | 0.72 |
| 1000 SW | 2.86 | 3.18 | 2.87 | 2.85 | 2.87 | 2.93 |
| BM | 5395 | 2793 | 3813 | 3400 | 4582 | 3926 |
| SY | 1150 | 547 | 794 | 751 | 956 | 830 |
| HI | 21.33 | 20.66 | 20.74 | 22.06 | 20.91 | 21.12 |
| OC | 50.46 | 48.08 | 49.62 | 49.09 | 49.37 | 49.30 |

Where, DF= Days to 50% flowering, DM=Days to 75% maturity, PH=Plant height (cm), LPBZ=Length of pods bearing zone (cm), PL= Pod length (cm), PW= Pod width (cm), PBP=Primary branch per plant, SBP=Secondary branch per plant, NPP= Number of pods per plant, BM= Biomass yield (kg/ha), 1000 SW=Thousand seed weight (g), SY= Seed yield (kg/ha), HI=Harvest index (%), and OC= Oil content (%)

According to the mean values of the clusters, selection of accession from Cluster I could be possible to obtain accession with the highest seed yield and other desirable traits. It is also suggested that accession from Cluster I and other Clusters be crossed to combine desirable traits and search for better-performing accession in subsequent segregating generations. The genotypes with the highest cluster mean value could be used as a parent in a future hybridization program for higher yield (Sultana *et al*., 2019).

From this study it can be concluded that 1) Five main components accounting to 76.3% total variation.2) for 64 sesame accessions, five distinct clusters were identified. 3) Highest inter-cluster distance between Cluster IV and Cluster V identified. 4) cluster I and cluster V had higher mean values over all mean values except for few traits as a result from those two clusters it could be possible to obtain accession with the highest seed yield and other desirable traits.

## Acknowledgement

I’m grateful to the Ethiopian Institute of Agricultural Research (EIAR) for allowing me to further my study and for providing me with the necessary financial assistance. My sincere thanks go to the Werer Agricultural Research Center staff for helping to manage field trials and set up the necessary infrastructure. For providing me with seeds from several sesame accessions and the pertinent data for my investigation, I am grateful to the Ethiopian Biodiversity Institute (EBI).

## REFERENCES

Ashim, C., Ghosh, P. D., and Sahu, P. K. (2013). Multivariate analysis of phenotypic diversity of landraces of rice of West Bengal. American Journal of Experimental Agriculture, 3(1): 110–123.

Bhatt, G. M. (1970). Multivariate analysis approach to selection of parents for hybridization aiming at yield improvement in self-pollinated crops. Australian Journal of Agricultural Research, 21(1): 1–7.

Brejda J.J., Moorman, T.B., Karlen, D.L. and Dao, T.H. (2000). Identification of regional soil quality factors and indicators I. Central and Southern High-Plains. Soil Science Society of America Journal, 64(6): 2115–2124.

CSA, R. (2019). The federal democratic republic of Ethiopia central statistical agency report on area and production of major. Statistical Bulletin.

CSA, R. (2020). The federal democratic republic of Ethiopia central statistical agency report on area and production of major. Statistical Bulletin.

CSA, R. (2021). The federal democratic republic of Ethiopia central statistical agency report on area and production of major. Statistical Bulletin.

Daniel E.G., and Parzies, H.K. (2011). Genetic variability among landraces of sesame in Ethiopia. African Crop Science Journal, 19(1):1–13.

Food and Agriculture Organization of the United Nations. (2020). Production of sesame. Top ten producer and exporter. FAOSTAT. Retrieved from https://www.fao.org/faostat/en

Getinet Alemawu, Geremew Terefe, Kasahun Zewdie, and Bulcha Weyassa. (1997). Lowland Oil Crops: A Three Decade Research Experiences in Ethiopia. Res. Report no. 31, Ethiopian Institute of Agricultural Research, Addis Ababa.

Gomez, K.A. and Gomez, A.A. (1984). Statistical procedures for agricultural research. John Wiley & Sons

IPGRI, N. (2004). Descriptors for sesame (Sesamumspp.). International Plant Genetic Resources Institute. Rome, Italy.

Morinaga, T., Fukushima, E., Kano, T., Maruyama, Y., and Yamasaki, Y. (1929). Chromosome numbers of cultivated plants II. Shokubutsugaku Zasshi, 43(515), 589–594.

Robbelen, G., Downey, R. K., & Ashri, A. (1989). Oil crops of the world: their breeding and utilization. McGraw-Hill New York P. 553

SAS Institute. (2014). SAS/STAT guide for personal computers, version 9.3 edition. AS Institute Inc., Cary, NC.

Shammoro MH, Kebede SA, and Gebremichael DE (2020). Genetic variability and multivariate analysis in indigenous and exotic sesame (*Sesamum indicum* L.) International Journal of Plant Breeding and Crop Science, 7(2): 814–823.

Sneath, P. H., & Sokal, R. R. (1973). Numerical taxonomy: the principles and practice of numerical classification.

Sultana, S., Mahmud, F., and Rahim, M. A. (2019). Genetic variability studies for selection of elite germplasm in sesame (Sesamum indicum L.). Agronomski glasnik: Glasilo Hrvatskog agronomskog društva, 81(2): 87–104.

Tesfaye, T., Tesfaye, K., Keneni, G., and Alemu, T. (2021). Morphological characteristics and genetic diversity of Ethiopian sesame genotypes. African Crop Science Journal, 29(1):59–76.

